# Variational quantum thermalization of environment-coupled DNA sequence biases

**DOI:** 10.64898/2026.10.30.763217

**Authors:** Hector Mejia-Diaz, Diego Santiago-Alarcon, Salvador E. Venegas-Andraca

**Affiliations:** Tecnologico de Monterrey, Escuela de Ingenieria y Ciencias, Monterrey, NL, 64849, Mexico; Department of Integrative Biology, University of South Florida, Tampa, Florida, 33620 USA

**Keywords:** quantum biology, variational quantum thermalizer, quantum evolution, point mutations, Bayesian optimization, quantum computing, Evolutionary Genomics

## Abstract

Understanding whether environmental conditions bias the statistical weight of specific genomic configurations requires a framework connecting molecular interactions, thermal fluctuations, and sequence-level observables. Here, we model DNA as an open quantum system coupled to an environmental thermal bath (represented as an external bosonic field) and parameterize the underlying molecular interaction energies (effective Hamiltonian terms) directly from prescribed nucleotide sequences. To evaluate sequence distributions at thermodynamic equilibrium, we compare two finite-temperature strategies at fixed inverse temperature (*β* = 1): classical Bayesian optimization targeting the exact thermodynamic equilibrium distribution (the Gibbs state), and quantum simulation using a Variational Quantum Thermalizer (VQT) implemented via a nine-layer parameterized quantum circuit. Both methods optimize the same interaction energy model and target-sequence objective across short oligomers (*N* = 2–5 base pairs), using 200 outer evaluations. Benchmarking against a uniform sequence distribution, exact-Gibbs optimization enhances target-sequence probabilities by factors of 9.11, 25.64, 54.51, and 159.87. The VQT approach similarly yields substantial target enrichment across all sequence lengths, achieving enhancement factors between 9.85 and 62.77. By evaluating the VQT-selected Hamiltonians against exact Gibbs calculations, we decouple optimization errors in parameter learning from approximation errors inherent to the variational quantum circuit. While concordance with the exact-Gibbs optimum is high in shorter oligomers, larger systems exhibit heightened sensitivity to variational thermal approximations. These results demonstrate that environment-induced sequence biases can be systematically captured within a variational quantum simulation framework, bridging biophysical models of DNA thermodynamics with quantum algorithms for finite-temperature state preparation. Crucially, the resulting equilibrium probabilities represent intrinsic thermo-dynamic biases rather than realized evolutionary substitution rates, which remain governed by kinetic activation barriers, replication fidelity, and enzymatic DNA repair.

## 1 Introduction

Since the middle of the twentieth century, evidence has accumulated showing that mutation is not necessarily uniform across sequence space and that mutational biases can influence evolutionary outcomes [1, 2]. Genomic studies have also described reproducible patterns of variation in parasites and bacteria [3, 4], while mutation-accumulation experiments show that base-substitution processes can differ across sequence contexts and organisms [5]. These observations motivate the search for physical descriptions able to connect environmental perturbations with sequence-dependent mutation probabilities.

Quantum biology provides examples in which quantum mechanical effects contribute to biological function, including energy transfer, enzymatic reactions, proton dynamics, olfaction, and magnetodetection [6, 7]. At the same time, quantum simulation was originally motivated by the difficulty of representing interacting quantum systems on classical computers [8]. If we can map out a biological process by looking at all the independent quantum states and vibrations that interact within it (i.e., interacting quantum degrees of freedom), using a quantum computer to iteratively simulate those states provides a powerful way to study how biological processes behave in the warm, chaotic environment of a real living organism. In evolutionary biology, quantum walks have also been proposed as a way of exploring genotype networks and evolutionary state spaces [9, 10].

Detailed microscopic models have addressed proton transfer and tunneling in individual DNA base pairs. Godbeer *et al*. [11] studied proton tunneling in adenine–thymine base pairs with a dissipative quantum model, while recent work by Slocombe *et al*. examined proton transfer using open-quantum-system dynamics and during DNA strand separation [12, 13], and Abrantes *et al*. [14] considered nuclear quantum effects and water-assisted proton transfer in methylated guanine–cytosine pairs. These approaches provide important microscopic information, but they also require a detailed description of the molecular geometry and of the specific reaction coordinate. A complementary approach is to describe the collective statistical behavior of a DNA–environment system without resolving each microscopic process independently.

In our previous work [15], we proposed an effective many-body energy model (Hamiltonian) in which the internal conformational and base-pairing degrees of freedom of DNA interact with a surrounding thermal bath (a bosonic environment) representing an external stressor. The model was designed so that environmental effects could be captured via coarse-grained coupling parameters, bypassing the need to model every molecular solvent–DNA interaction in full microscopic detail. We showed that physically plausible parameter ranges successfully reproduce biologically relevant energy scales (e.g., the thermodynamic energies governing hydrogen bonding and base stacking) and that coupling to this environment directly alters local thermal fluctuations. The free parameters in that Hamiltonian, however, were constrained solely by theoretical thermodynamic arguments rather than learned directly from target nucleotide sequences.

Here, we take the next step and optimize the effective Hamiltonian (i.e., the molecular energy operator specifying the energetic costs of nucleotide configurations, base stacking interactions, and environmental coupling) using prescribed nucleotide sequences. The central object is the finite-temperature Gibbs state:

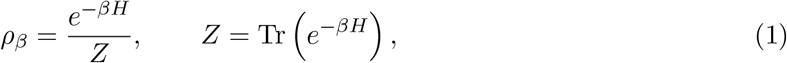

which provides the exact thermodynamic equilibrium distribution for a fixed Hamiltonian. In this formulation, *β* = 1*/*(*k*_*B*_*T* ) is the inverse temperature (where *k*_*B*_ is the Boltzmann constant and *T* is temperature), parameterizing the intensity of thermal fluctuations relative to molecular interaction energies: as *β* increases (lower temperature), the system concentrates in the lowest-energy states, while at smaller *β* (higher temperature), thermal noise distributes probability more uniformly across sequences. Equation (1) is the quantum analogue of the classical Boltzmann distribution: the exponential term *e*^*−βH*^ weights each configuration by its thermodynamic stability, while the partition function *Z* = Tr(*e*^*−βH*^) acts as the normalization factor summing over all accessible states so that total probability equals 1.

Because the total system includes both DNA and the surrounding bath, we trace out the environmental degree of freedom—the mathematical equivalent of integrating out or marginalizing over the external solvent and stressor states. The diagonal elements of the resulting reduced DNA density matrix define a classical probability distribution over discrete nucleotide strings (*A, C, G, T* ). We optimize the Hamiltonian parameters so that a prescribed target sequence occupies a free-energy minimum and receives enhanced statistical weight.

We compare two strategies. The first uses Bayesian optimization together with direct evaluation of Eq. (1); throughout the paper we refer to this as exact-Gibbs BO. The second uses the Variational Quantum Thermalizer (VQT) [16, 17], which approximates a thermal state by combining a classical probability distribution with a parameterized quantum circuit. The simulations were implemented in PennyLane [18]. Here, VQT provides a finite-temperature variational formulation in which thermal-state preparation is expressed through a parameterized quantum circuit and a classical probability distribution.

Thermodynamic equilibrium probabilities and experimentally measured mutation rates describe distinct biological quantities. A thermal Gibbs distribution reflects the static thermodynamic stability and physical accessibility of a base configuration or change. In contrast, realized evolutionary mutation rates, measured per base pair per generation across benchmark organisms such as *Drosophila*, humans, and *Caenorhabditis elegans* [19, 20, 5], are governed by extensive downstream biological filtering. In living systems, DNA damage processing is actively mediated by enzymatic repair pathways, such as nucleotide excision repair resolving ultraviolet-induced alterations [21, 22], while in vivo replication fidelity depends on DNA polymerase selectivity, 3^*′*^ to 5^*′*^ exonucleolytic proofreading, and post-replicative mismatch repair [23, 24]. Therefore, translating an equilibrium thermodynamic probability into a dynamic substitution rate requires an independently parameterized kinetic model that accounts for transition-state activation barriers, replication cycles, and enzymatic repair rates. For this reason, the present study focuses strictly on relative sequence-probability enhancement (thermodynamic enrichment over a uniform baseline) rather than on a direct conversion to realized mutation rates.

Finally, environmental effects on DNA are not restricted to radiation. Electrostatic and ionic conditions can modify DNA free-energy landscapes [25], and experimentally measured nearest-neighbor base-pair free energies depend on salt concentration and sequence composition [26]. These observations motivate the broader environmental-stressor framework of the model, in which external conditions are represented through effective couplings to the DNA degrees of freedom.

The purpose of the present comparison is therefore twofold: first, to determine whether the effective DNA–environment Hamiltonian (the molecular interaction energy model) can be parameterized so that prescribed mutation target sequences (such as known motifs prone to changes or mutational hotspots) acquire enhanced thermodynamic weight at equilibrium; and second, to test whether this same finite-temperature objective can be accurately prepared on quantum hardware using a Variational Quantum Thermalizer (VQT) circuit. Exact Gibbs calculations provide the classical analytical ground truth, while the VQT results establish a quantum algorithmic framework for sampling the resulting environment-dependent sequence biases.

## 2 Methods

### 2.1 Sequence encoding and benchmark targets

Each nucleotide is represented by two qubits according to

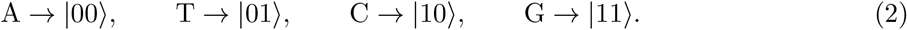

We considered target subsequences of the string GGTATTAATG with lengths *N* = 2, 3, 4, and 5, namely GG, GGT, GGTA, and GGTAT. The comparison was truncated at *N* = 5 because the dense exact-Gibbs diagnostic used for the matched analysis exceeded the available 16 GB memory at *N* = 6.

For the numerical benchmark, the external environmental stressor is modeled using a single vibrational bath mode truncated to a two-level system (*D* = 2, corresponding to an unperturbed vs. excited stressor state) and represented by one additional qubit. Because each nucleotide position (*A, C, G, T* ) requires two qubits to encode its four discrete states (2^2^ = 4), an oligomer of length *N* base pairs occupies 2*N* qubits; including the environmental qubit yields a total quantum register of 2*N* + 1 qubits with a state-space dimension of 2^2*N*+1^ (representing 2 × 4^*N*^ joint sequence– environment configurations). The inverse temperature was fixed at *β* = 1 (in standard reduced simulation units where thermal energy *k*_*B*_*T* = 1) to establish a consistent thermal baseline for the matched comparison.

### 2.2 Effective Hamiltonian

We retain the local and environmental interaction structure introduced in our previous work [15]. In the original formulation,

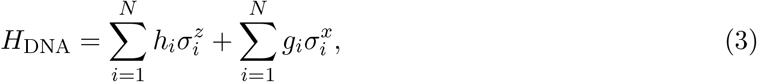

and

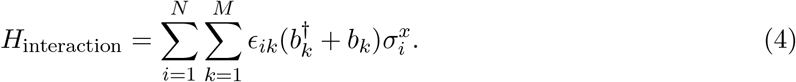

Because the present nucleotide encoding uses two qubits per nucleotide, the same local structure is extended to both encoded qubit degrees of freedom. For a single truncated environmental mode, the implemented Hamiltonian is

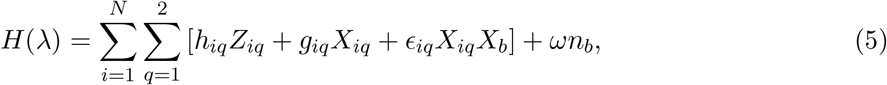

where *b* denotes the bosonic bath qubit, *Z*_*b*_ is the Pauli-*Z* operator acting on that bath qubit, and

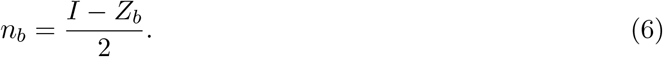

Here *n*_*b*_ is the bath occupation-number operator. For the *D* = 2 truncation, *b*^*†*^ + *b* is represented by *X*_*b*_, the Pauli-*X* operator acting on the bath qubit. The coefficients *h*_*iq*_, *g*_*iq*_, and *ϵ*_*iq*_ describe the nucleotide and DNA–environment interaction terms, while *ω* is the characteristic energy of the environmental mode. The parameter vector is therefore

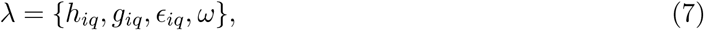

with 6*N* + 1 free Hamiltonian parameters (two encoded qubits per nucleotide, with three coefficients *h*_*iq*_, *g*_*iq*_, and *ϵ*_*iq*_ for each qubit, plus the single environmental parameter *ω*). We use the bounds

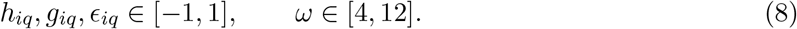

The model represents an effective thermal Hamiltonian rather than as a microscopic description of individual proton-transfer events.

### 2.3 Exact Gibbs reference and Bayesian optimization

For a given set of interaction parameters *λ* (the tunable base-pairing, stacking, and environmental couplings), the dense Hamiltonian matrix is constructed and the exact Gibbs state *ρ*_*β*_ is obtained from Eq. (1). To isolate the sequence probabilities, the environmental bath qubit is traced out (i.e., marginalizing over all external stressor states), yielding the reduced DNA density matrix:

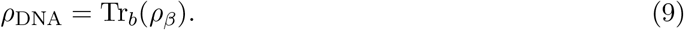

For a target sequence |*ψ*_target_⟩ (the specific nucleotide sequence or genotype of interest), the predictive quantity is its thermodynamic equilibrium probability:

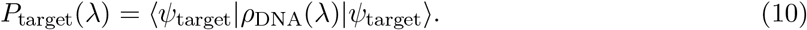

which isolates the diagonal entry of *ρ*_DNA_ corresponding to the target sequence.

The outer optimization algorithm searches parameter space *λ* by minimizing the negative log probability:

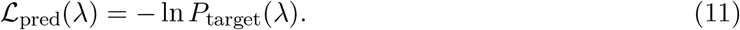

which is mathematically equivalent to maximizing the statistical weight (likelihood) of observing that target genotype at equilibrium.

We used Gaussian-process Bayesian optimization with expected improvement, 200 total objective evaluations, and 30 initial points. These outer-search settings were also used in the VQT comparison.

To assess the global energetics of the system, we also computed the ensemble mean energy:

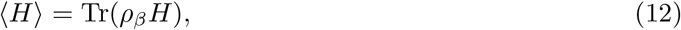

and the energy variance:

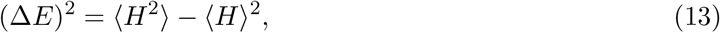

which characterizes the magnitude of thermal energy fluctuations, measuring how broadly thermal noise disperses the system across different conformational energy states around the mean.

### 2.4 Variational Quantum Thermalizer

The Variational Quantum Thermalizer (VQT) variationally minimizes the Helmholtz free energy [16, 17]:

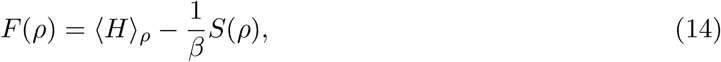

which formalizes the fundamental thermodynamic trade-off between minimizing internal molecular energy ⟨*H*⟩_*ρ*_ and maximizing state entropy *S*(*ρ*)—the quantum equivalent of the classical thermodynamic relation *F* = *E* − *TS* (noting that *T* = 1*/*(*k*_*B*_*β*)).

Here, *S*(*ρ*) = − Tr(*ρ* log *ρ*) is the von Neumann entropy, which serves as the quantum counterpart of Shannon entropy, quantifying the statistical disorder or diversity across accessible sequence states in the thermal ensemble. Because a finite-temperature thermal state is an ensemble (a mixed state with classical uncertainty) rather than a single pure quantum state, the algorithm introduces entropy by sampling initially from an independent Bernoulli distribution over computational-basis states—analogous to a site-independent allele-frequency model in population genetics:

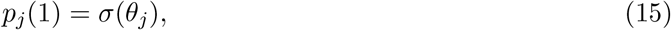

where *σ*(*z*) = 1*/*(1 + *e*^*−z*^) is the standard logistic sigmoid function parameterized by trainable classical parameters *θ*_*j*_. For each sampled bitstring **s** (representing an initial classical sequence configuration), a parameterized quantum circuit applies a unitary transformation *U*_*ϕ*_ that introduces inter-qubit correlations and quantum superpositions, yielding the variational mixed state:

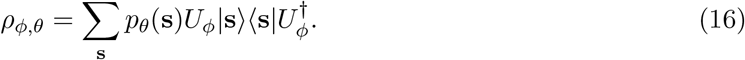

The complete variational circuit and the nested optimization strategy are summarized in Fig. 1. This nested architecture decouples the physical simulation from the parameter-fitting task:

**Figure 1:**
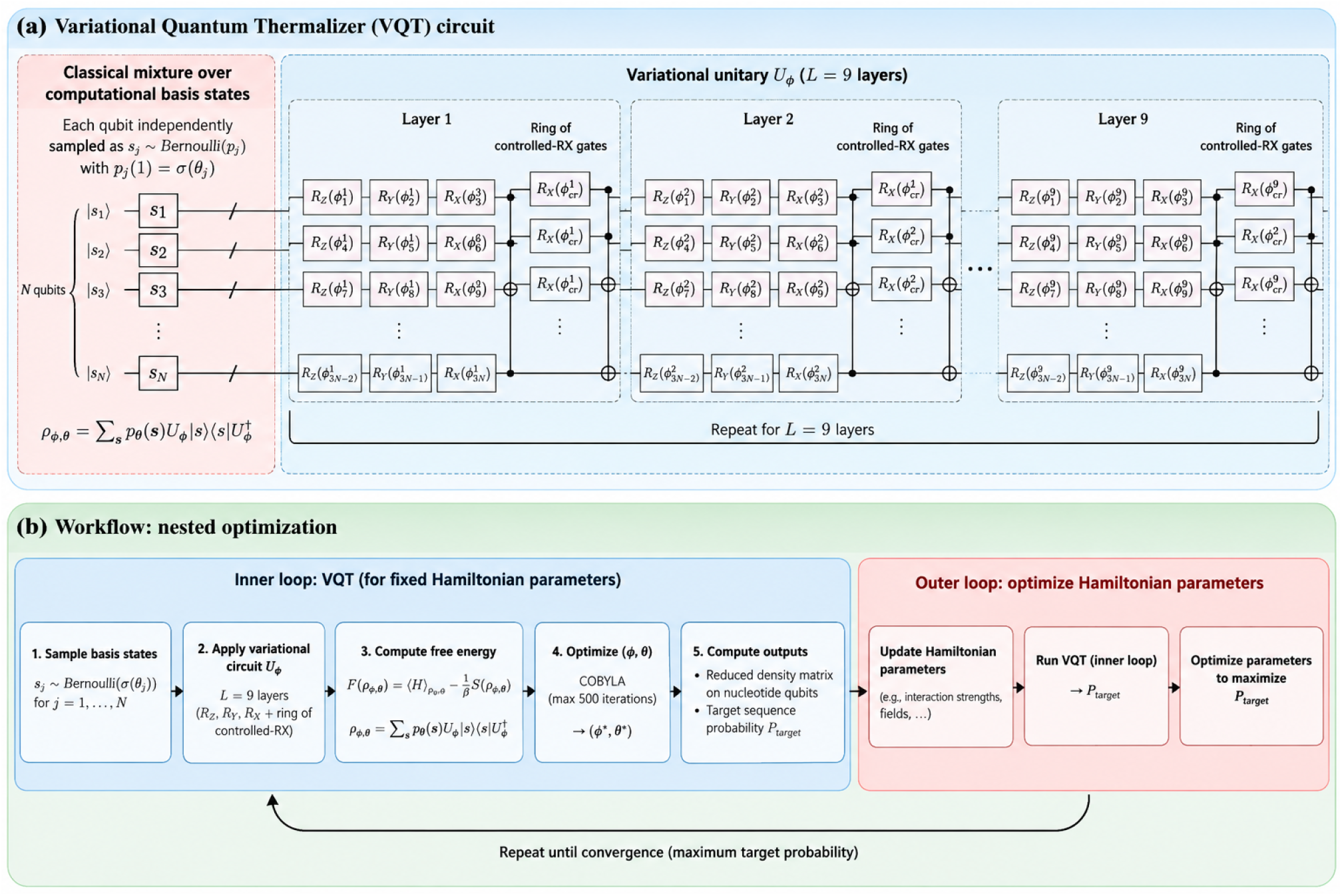
Variational Quantum Thermalizer circuit and nested optimization workflow. **(a)** The VQT constructs a mixed thermal-state approximation from a classical product distribution over computational-basis states and a parameterized quantum circuit. Each sampled basis state is evolved by a variational unitary *U*_ϕ_ composed of *L* = 9 repeated layers of single-qubit *R*_Z_, *R*_Y_, and *R*_X_ rotations followed by a ring of controlled-*R*_X_ entangling gates. The resulting mixed state is 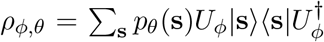. **(b)** Nested optimization procedure used for prediction. For fixed Hamiltonian parameters, the inner VQT loop minimizes the free energy by optimizing the classical distribution parameters and circuit parameters. The resulting reduced density matrix of the nucleotide register is used to evaluate the target-sequence probability *P*_target_. This probability is then passed to the outer optimization loop, which updates the Hamiltonian parameters to maximize *P*_target_. The procedure is repeated until the outer optimization converges.

- **Inner loop (Physics simulation):** For a fixed set of Hamiltonian parameters, the inner VQT loop optimizes the classical distribution parameters *θ* and circuit parameters *ϕ* to minimize the free energy *F* (*ρ*), thereby preparing the approximate thermodynamic equilibrium (Gibbs state) for that specific energy model.
- **Outer loop (Parameter learning):** The resulting target-sequence probability *P*_target_ is evaluated from the reduced density matrix of the nucleotide register, and the outer loop updates the Hamiltonian interaction parameters to maximize this target probability.

The circuit contains nine layers (*L* = 9). Each layer applies single-qubit rotations (*R*_*Z*_, *R*_*Y*_, *R*_*X*_) on every qubit to adjust individual state amplitudes, followed by two-qubit controlled-*R*_*X*_ gates arranged in a ring topology to generate inter-site coupling and entanglement. The inner free-energy optimization is performed using the gradient-free COBYLA algorithm. During the outer Hamiltonian search, we used 120 inner function evaluations, 30 energy samples, and 300 samples for the target-probability estimate. Variational parameters were warm-started between successive outer evaluations. After the 200-call outer search, the selected Hamiltonian was refit using 600 inner evaluations, 100 energy samples, and 3,000 samples for the final predictive estimate.

The VQT outer objective is the same negative log-likelihood defined in Eq. (11). Therefore, exact-Gibbs Bayesian optimization and VQT optimize the exact same Hamiltonian family, parameter bounds, inverse temperature (*β* = 1), target sequences, and number of outer calls (200 evaluations). They differ solely in the computational engine used to obtain the thermal-state probabilities: exact classical matrix exponentiation versus variational simulation on a quantum circuit.

### 2.5 Exact-Gibbs diagnostic at the VQT Hamiltonian

To distinguish errors in the Hamiltonian search from errors in the variational thermal-state approximation, we evaluate the exact Gibbs state at the Hamiltonian selected by the VQT outer loop. We denote this quantity by

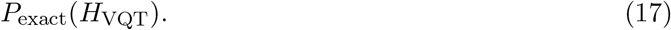

Comparing *P*_BO_, *P*_exact_(*H*_VQT_), and *P*_VQT_ separates two effects: whether the outer search found a favorable Hamiltonian, and whether VQT accurately represents the thermal state of that Hamiltonian.

### 2.6 Environmental-coupling sensitivity

To assess the specific role of the environmental interaction independently of the remaining Hamiltonian parameters, we evaluate the exact Gibbs target probability after setting all optimized environmental couplings to zero (*ϵ* = 0) while keeping the intrinsic DNA and bare stressor parameters—the single-site nucleotide biases (*h*_*iq*_), the inter-site stacking and pairing interactions (*g*_*iq*_), and the characteristic stressor energy (*ω*)—fixed. This serves as an in silico ablation control, isolating whether target-sequence enrichment is actively driven by external stressor coupling rather than intrinsic DNA stability alone.

We compare the fully coupled target probability:

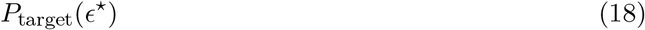

with the uncoupled baseline:

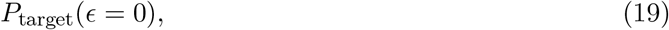

and define the environmental sensitivity ratio (fold-enrichment):

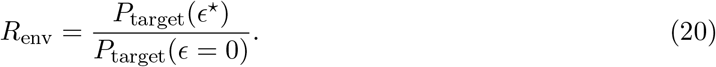

Values above or below unity indicate that the selected environmental coupling respectively increases (*R*_env_ *>* 1) or decreases (*R*_env_ *<* 1) the target-sequence probability relative to the corresponding uncoupled Hamiltonian. In evolutionary terms, *R*_env_ *>* 1 demonstrates that interaction with the environmental stressor actively enhances the thermodynamic accessibility of that specific sequence configuration beyond what intrinsic base-stacking and pairing energetics dictate in isolation.

## 3 Results

### 3.1 Exact-Gibbs Bayesian optimization

Because the number of possible nucleotide strings grows as 4^*N*^, the probability of an individual sequence decreases rapidly with *N* . We therefore compare *P*_target_ with the uniform reference

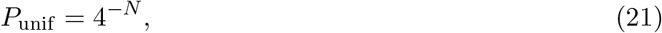

and define the enhancement

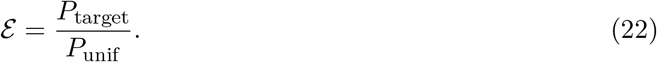

An enhancement larger than one means that the optimized thermal state assigns the target more probability than an unbiased distribution over nucleotide strings.

Exact-Gibbs BO gives target probabilities of 0.56950, 0.40061, 0.21291, and 0.15612 for *N* = 2– 5, respectively. The corresponding uniform probabilities are 0.0625, 0.015625, 0.00390625, and 0.0009765625. Although the absolute target probability decreases as the sequence length grows, the relative enhancement increases strongly, from 9.11 at *N* = 2 to 159.87 at *N* = 5. See Fig. 2 and Table 1.

**Table 1:** Matched predictive results at *β* = 1. The exact-Gibbs diagnostic at the VQT-selected Hamiltonian separates Hamiltonian-search quality from variational thermal-state error.

| *N* | Target | *P*<sub>BO</sub> | *P*<sub>exact</sub>(*H*<sub>VQT</sub>) | *P*<sub>VQT</sub> | ℰ<sub>VQT</sub> |
|---|---|---|---|---|---|
| 2 | GG | 0.56950 | 0.53447 | 0.72532 | 11.61 |
| 3 | GGT | 0.40061 | 0.28414 | 0.55758 | 35.69 |
| 4 | GGTA | 0.21291 | 0.04615 | 0.03847 | 9.85 |
| 5 | GGTAT | 0.15612 | 0.02936 | 0.06130 | 62.77 |

**Table 2:**
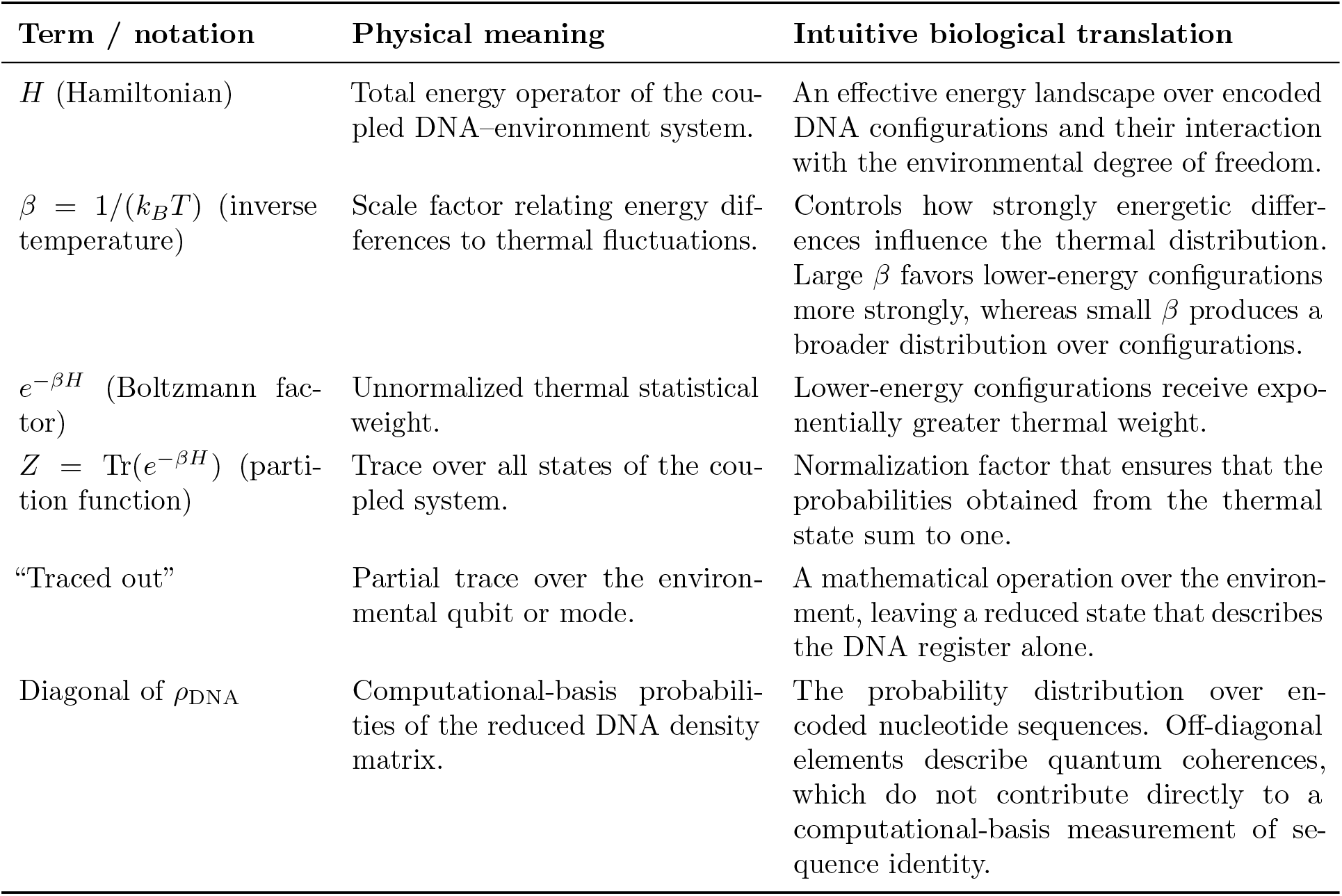
Translation of the main statistical-mechanical quantities into biologically intuitive language.

| **Term / notation** | **Physical meaning** | **Intuitive biological translation** |
|---|---|---|
| *H* (Hamiltonian) | Total energy operator of the cou-pled DNA–environment system. | An effective energy landscape over encoded DNA configurations and their interaction with the environmental degree of freedom. |
| *β* = 1/(*k*<sub>*B*</sub>*T*) (inverse temperature) | Scale factor relating energy dif-ferences to thermal fluctuations. | Controls how strongly energetic differ-ences influence the thermal distribution. Large *β* favors lower-energy configurations more strongly, whereas small *β* produces a broader distribution over configurations. |
| *e*<sup>−*βH*</sup> (Boltzmann fac-tor) | Unnormalized thermal statistical weight. | Lower-energy configurations receive expo-nentially greater thermal weight. |
| *Z* = Tr(*e*<sup>−*βH*</sup>) (parti-tion function) | Trace over all states of the cou-pled system. | Normalization factor that ensures that the probabilities obtained from the thermal state sum to one. |
| “Traced out” | Partial trace over the environ-mental qubit or mode. | A mathematical operation over the environ-ment, leaving a reduced state that describes the DNA register alone. |
| Diagonal of *ρ*<sub>DNA</sub> | Computational-basis probabili-ties of the reduced DNA density matrix. | The probability distribution over en-coded nucleotide sequences. Off-diagonal elements describe quantum coherences, which do not contribute directly to a computational-basis measurement of se-quence identity. |

**Figure 2:**
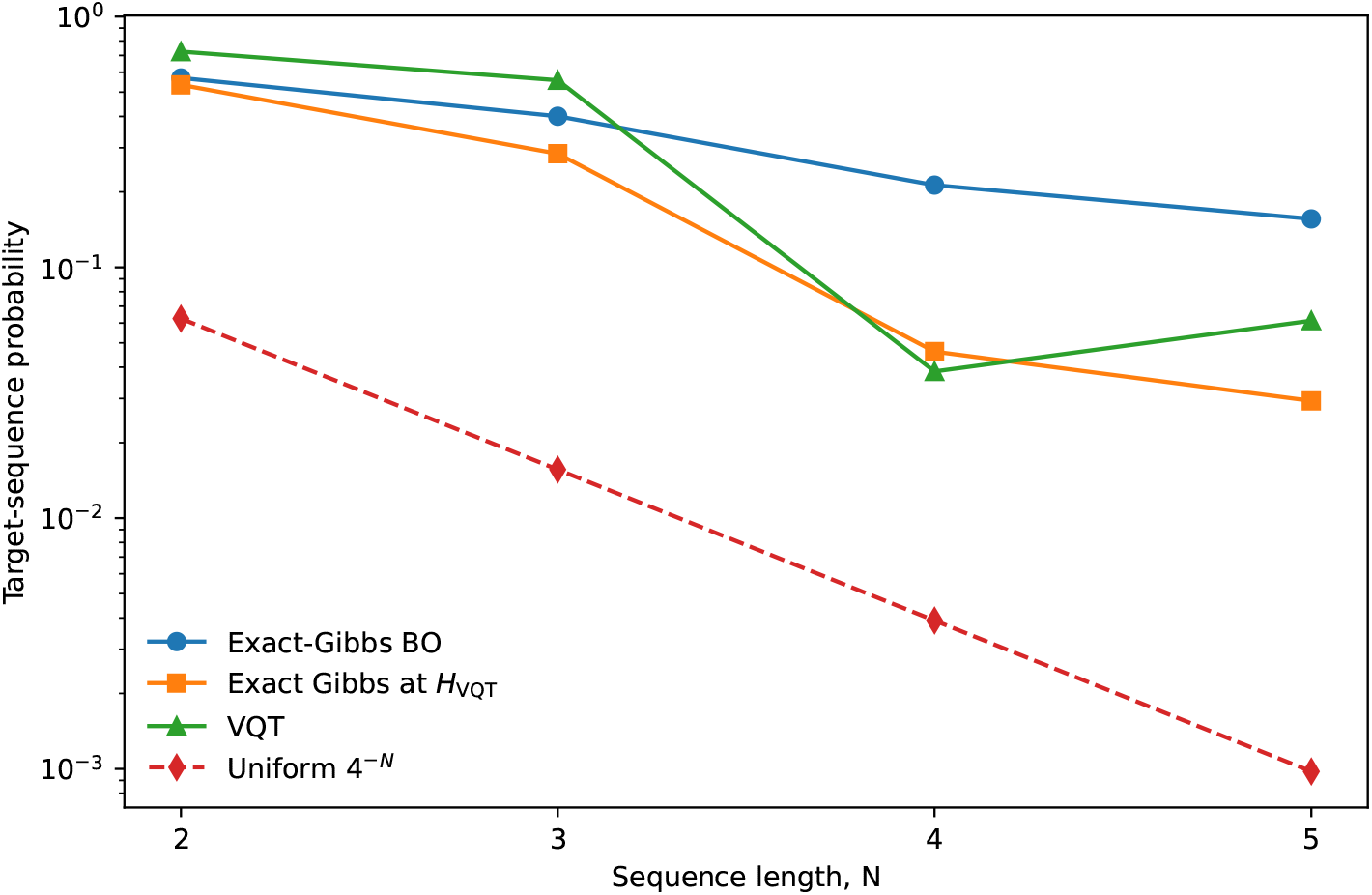
Target-sequence probabilities for exact-Gibbs BO, the exact Gibbs state evaluated at the Hamiltonian selected by the VQT-guided search, and the final VQT state. The uniform reference 4^*−N*^ is included for comparison.

For example, at *N* = 5 there are 4^5^ = 1024 possible nucleotide strings. A uniform distribution gives the target GGTAT a probability 1*/*1024 ≈ 9.77×10^*−*4^, whereas the optimized exact Gibbs state gives *P*_target_ = 0.15612. The target therefore carries approximately 159.87 times more statistical weight than under a uniform distribution, see Fig. 3. This result should be interpreted as a learned sequence-selective thermal bias rather than as a mutation rate.

**Figure 3:**
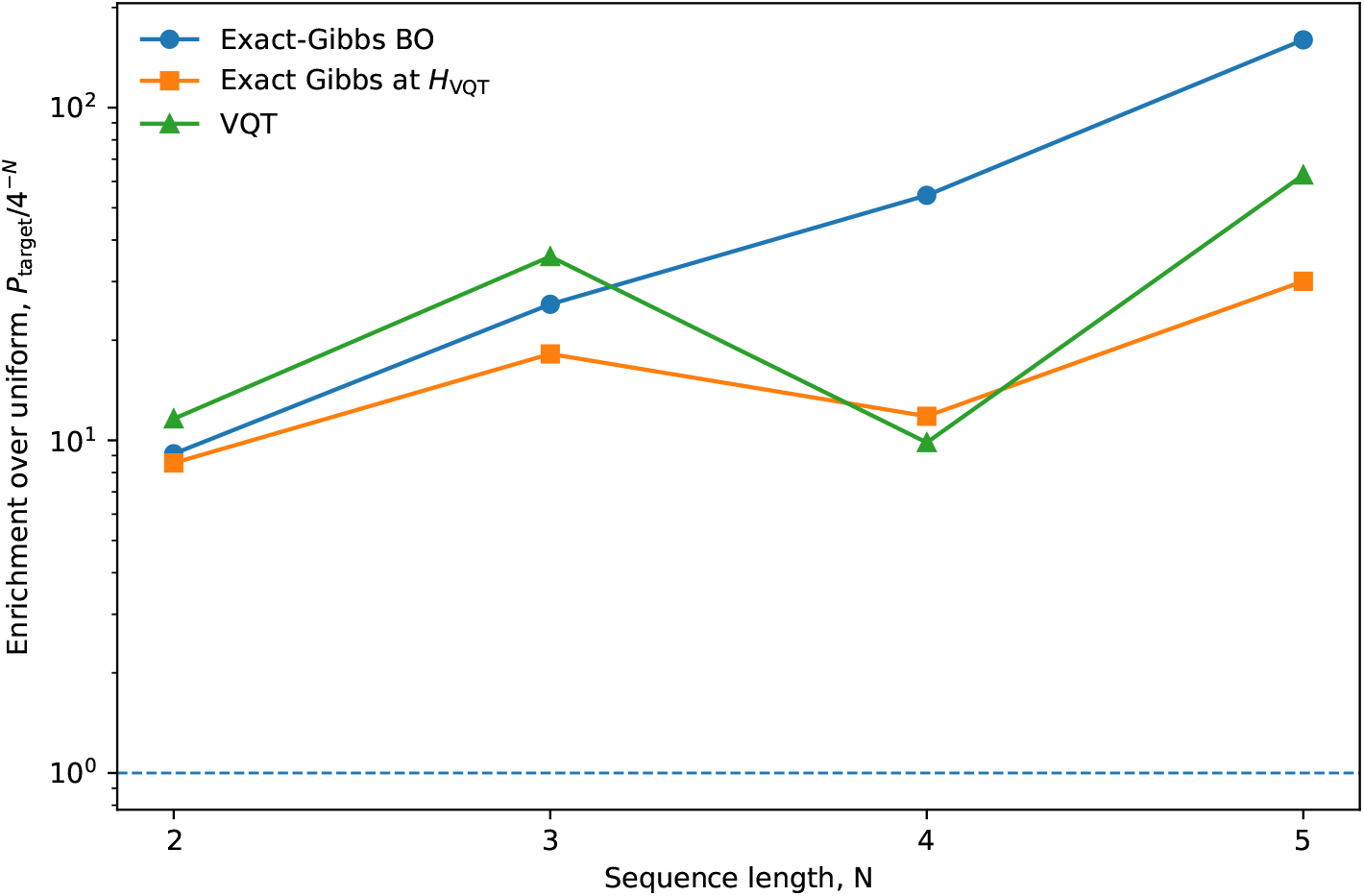
Enhancement of the prescribed target sequence relative to the uniform reference 4^*−N*^ . Exact-Gibbs BO produces increasingly strong relative enrichment as the sequence space grows. VQT retains target enrichment above the uniform background for all sequence lengths considered.

### 3.2 Predictive VQT and comparison with the exact Gibbs state

VQT also produces a target probability above the uniform reference for every system considered. The final VQT probabilities are 0.72532, 0.55758, 0.03847, and 0.06130 for *N* = 2–5, corresponding to enhancement factors of 11.61, 35.69, 9.85, and 62.77, respectively. See Figure 2 and Table 1.

The comparison in Fig. 2 shows that the origin of the difference between BO and VQT varies across system size. At *N* = 2, the Hamiltonian identified by the VQT-guided search gives *P*_exact_(*H*_VQT_) = 0.53447, which is close to the BO value 0.56950. Thus, for the smallest system the VQT-guided outer search identifies a Hamiltonian with a target probability close to the exact-Gibbs BO solution (Table 1).

At *N* = 3, the VQT-selected Hamiltonian also retains substantial exact-Gibbs target weight, with *P*_exact_(*H*_VQT_) = 0.28414 compared with *P*_BO_ = 0.40061. For *N* = 4 and *N* = 5, the corresponding values fall to 0.04615 and 0.02936, respectively, indicating that the VQT-guided outer search increasingly samples a different effective optimization landscape from the one obtained using exact Gibbs probabilities.

The final target probability estimated by the quantum circuit (*P*_VQT_) does not systematically overestimate or underestimate the exact thermodynamic equilibrium probability (*P*_exact_(*H*_VQT_)) for the same underlying Hamiltonian. For sequence lengths *N* = 2, 3, and 5 base pairs, the VQT circuit yields a slightly higher target probability than the exact Gibbs state (*P*_VQT_ *> P*_exact_), whereas at *N* = 4 it yields a lower probability (*P*_VQT_ *< P*_exact_). This alternating sign demonstrates that the variational thermal approximation does not introduce a simple, one-directional biological bias—such as systematically exaggerating target accessibility (acting as an artificial mutational hotspot) or systematically over-dispersing probability across non-target sequences (acting as an artificial entropic noise floor). Instead, deviations reflect finite-circuit expressivity and numerical optimization variance around the true equilibrium landscape (Fig. 4).

**Figure 4:**
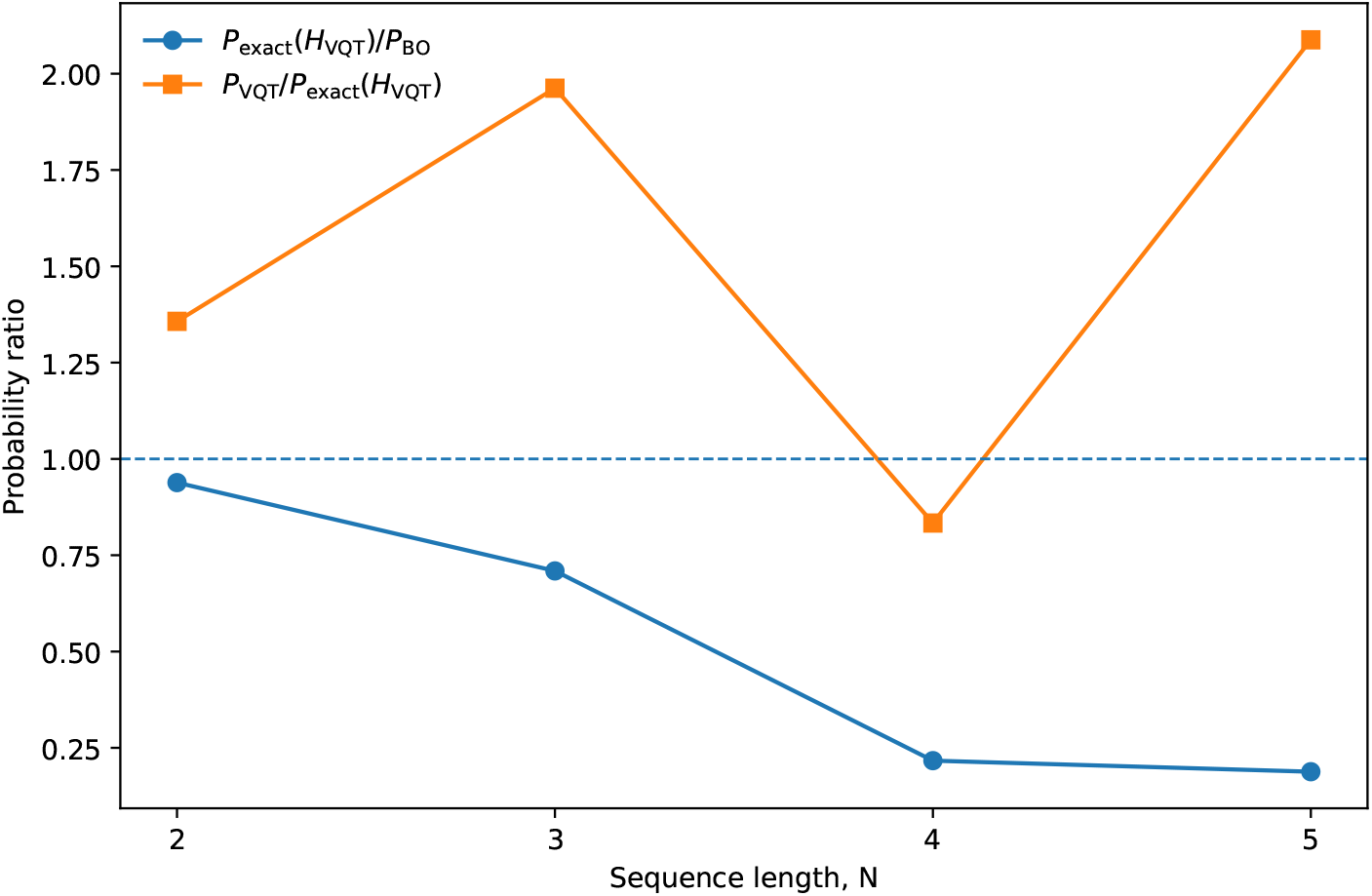
Diagnostic ratios separating Hamiltonian-search quality from variational thermal-state approximation. The ratio *P*_exact_(*H*_VQT_)*/P*_BO_ measures how favorable the VQT-selected Hamiltonian is under exact Gibbs evaluation, while *P*_VQT_*/P*_exact_(*H*_VQT_) compares the final variational probability with the exact Gibbs probability at the same Hamiltonian.

### 3.3 Environmental coupling and sequence probabilities

Changing the environmental interaction while holding the remaining Hamiltonian parameters fixed can either increase or decrease the target-sequence probability.

For the BO-selected Hamiltonians, *R*_env_ is approximately 0.958, 0.954, 1.002, and 0.728 for *N* = 2–5. For the VQT-selected Hamiltonians evaluated exactly, the corresponding values are approximately 0.974, 0.900, 0.935, and 0.929. Thus, in the present optimized parameter sets, the environmental interaction can either slightly favor or suppress the target sequence. The biologically relevant interpretation is therefore that the environment reshapes the effective sequence landscape (Fig. 5).

**Figure 5:**
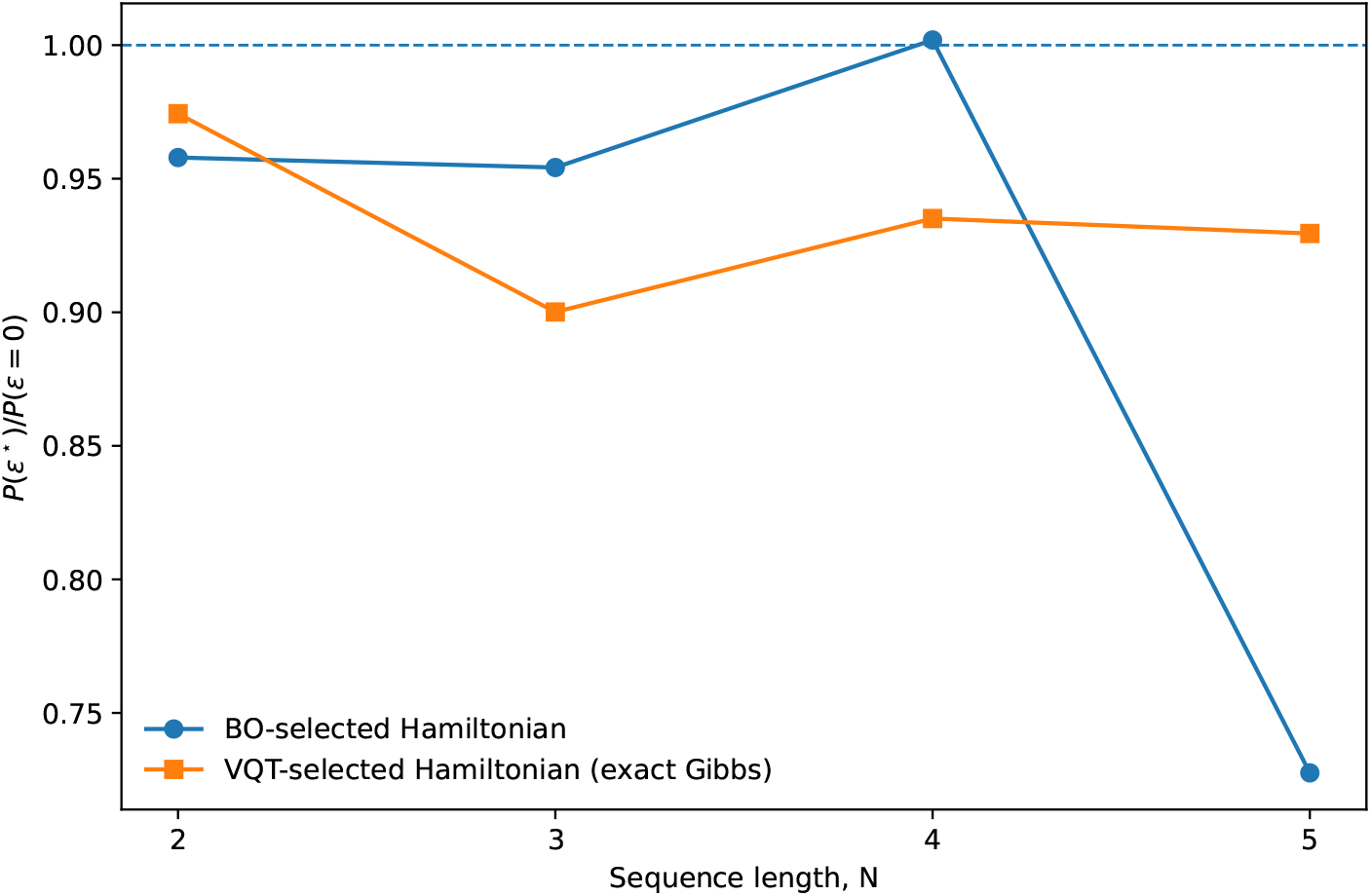
Environmental-coupling sensitivity. The ratio *P* (*ϵ*^⋆^)*/P* (*ϵ* = 0) compares the target probability at the selected environmental couplings with the probability obtained after setting the environmental interaction to zero while holding the remaining Hamiltonian parameters fixed. Values above or below unity indicate relative enhancement or suppression, respectively.

### 3.4 Thermal fluctuations

The exact-Gibbs Bayesian optimization states also provide the thermal energy variance (Δ*E*)^2^ of the optimized Hamiltonians (Eq. (13)). At inverse temperature *β* = 1, (Δ*E*)^2^ takes values of approximately 1.64, 2.46, 3.28, and 4.05 for sequence lengths *N* = 2–5 base pairs, respectively. This steady, near-linear increase (∼ 0.82 per nucleotide) is consistent with an extensive physical property, where each additional base pair introduces independent vibrational and conformational degrees of freedom to the thermal ensemble. Biologically, we interpret these values as localized thermal fluctuations—the routine “breathing” and base-stacking noise of a finite molecular system— rather than signatures of a cooperative thermodynamic phase transition, such as duplex melting or cooperative structural collapse.

### 3.5 Computational cost

The computational cost differs strongly between the two approaches when both are executed classically. Exact-Gibbs BO required approximately 0.49, 0.78, 0.91, and 1.16 h for *N* = 2–5, respectively. The corresponding VQT calculations required approximately 1.64, 3.26, 3.75, and 7.25 h.

These timings reflect the present classical implementations of both methods. The VQT circuits are simulated with a classical state-vector backend, and the nested variational procedure introduces substantial overhead. The relevance of VQT in the present work is that the same DNA–environment thermal problem is expressed as a parameterized quantum circuit for finite-temperature state preparation. In contrast, direct dense construction of the exact Gibbs state requires explicit manipulation of exponentially growing matrices. In our implementation, the dense exact-Gibbs diagnostic at *N* = 6 exceeded the available 16 GB memory, so the matched comparison was restricted to *N* ≤ 5. The measured wall-clock scaling is shown in Fig. 6.

**Figure 6:**
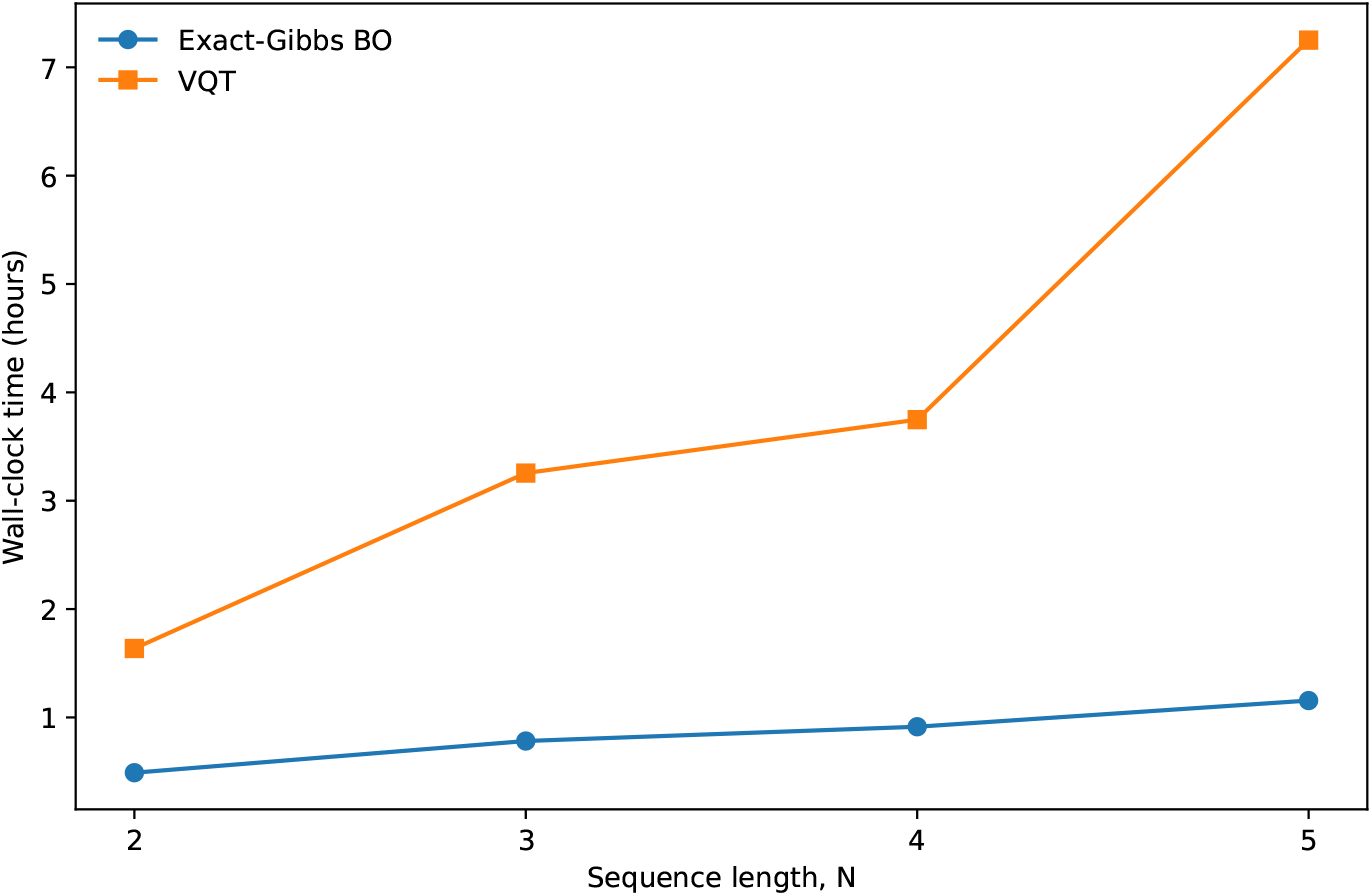
Measured wall-clock time for the matched calculations. VQT is more expensive in the present classical simulation because each Hamiltonian evaluation contains an inner variational thermalization and repeated circuit sampling.

## 4 Discussion

The exact-Gibbs calculations establish a rigorous quantitative baseline for the effective DNA– environment Hamiltonian. Across every oligomer length evaluated, Bayesian optimization concentrates statistical weight onto the prescribed sequence well above the uniform null expectation, with the fold-enhancement increasing monotonically from 9.11 at *N* = 2 to 159.87 at *N* = 5 base pairs. While the absolute probability of observing any single sequence inevitably declines as sequence space expands, it decays substantially slower than the unbiased combinatorial background rate of 4^*−N*^ . The learned Hamiltonian family therefore yields increasingly pronounced sequence selectivity as oligomer length grows, demonstrating its capacity to encode strong, sequence-dependent thermodynamic biases.

Biologically, this scaling behavior delivers an essential insight: environmental coupling provides a direct biophysical mechanism for breaking the assumed isotropy of sequence space. Evolutionary models frequently treat single-step mutations across genotype networks as equiprobable or governed by flat substitution matrices [27]. Our results show that physical coupling to an external stressor can compress high-dimensional combinatorial space, focusing thermodynamic probability onto specific sequence motifs by more than two orders of magnitude. This demonstrates that the raw mutational input available to evolution is not an unbiased random walk [28], but can be pre-channeled by environmental biophysics into localized hotspots before downstream kinetic barriers, enzymatic repair pathways, and natural selection intervene.

The VQT results demonstrate that the sequence-bias objective can be successfully mapped onto a finite-temperature variational quantum algorithm of the type introduced in Refs. [16, 17]. Across all oligomer lengths evaluated, the target genotype remains substantially enriched above the uniform null baseline. Crucially, evaluating the exact Gibbs distribution on the VQT-derived Hamiltonian (*H*_VQT_) provides an essential diagnostic by decoupling two distinct sources of error that would otherwise remain confounded: the parameter-learning efficiency of the outer search versus the physical approximation fidelity of the variational quantum circuit. In evolutionary modeling terms, this cleanly separates model inference error (identifying the correct biophysical energy landscape) from state-sampling error (how accurately the algorithm explores that landscape at finite temperature).

For short sequences (*N* = 2 base pairs), the VQT-guided outer search recovers an energy parameterization nearly indistinguishable from the exact classical Bayesian optimization optimum, and at *N* = 3 it preserves a substantial fraction of the exact-Gibbs target probability. For larger sequence spaces (*N* = 4 and *N* = 5), the Hamiltonians selected by the VQT search yield lower target probabilities when benchmarked against exact Gibbs calculations. This discrepancy reflects the inherent trade-offs of embedding an approximate thermal solver within a stochastic optimization loop: finite measurement sampling (analogous to genetic drift or finite population sampling noise), constrained inner iteration budgets, and the limits of a nine-layer circuit distorts the effective landscape traversed by the optimizer.

Beyond benchmarking quantum circuit fidelity, these results demonstrate a viable computational path toward solving the problem of environmental mutagenesis. In evolutionary genomics, we routinely observe non-random sequence patterns—such as mutational signatures in cancer genomes, CpG depletion, or codon-usage biases under environmental stress—and seek to infer the underlying biophysical constraints that generated them. Classical calculation of the exact thermodynamic partition function (*Z* = ∑*e*^*−βE*^) scales exponentially (4^*N*^ ), becoming computationally intractable beyond short oligomers and forcing evolutionary models to fall back on phenomenological, context-independent substitution matrices.

By demonstrating that a parameterized quantum circuit can recover sequence-enriching Hamil-tonians across expanding state spaces, this framework establishes proof-of-concept for reverse-engineering environmental mutational biases from first principles. It shows that quantum simulation can capture how external physical stressors shape the raw mutational spectrum—the primary variational repertoire that determines which evolutionary pathways natural selection can access.

The *N* = 4 base-pair oligomer result is particularly informative in this respect. At this sequence length—where sequence space expands to 4^4^ = 256 discrete genotypes—the final probability estimated by the VQT circuit (*P*_VQT_ = 0.03847) remains reasonably close to the exact thermodynamic equilibrium probability evaluated on that same Hamiltonian (*P*_exact_(*H*_VQT_) = 0.04615), even though both remain below the global optimum reached by direct classical Bayesian optimization. This agreement indicates that the primary bottleneck at this scale is the parameter set selected by the VQT-guided outer search, rather than an inability of the quantum circuit to accurately represent the physical thermal distribution of the chosen Hamiltonian.

This diagnosis delivers an encouraging conclusion for evolutionary genetics: the quantum hardware can faithfully reproduce the equilibrium landscape of a multi-site genotype network without breakdown. In an *N* = 4 system (encoded on a 9-qubit register, 2*N* +1 qubits), the circuit successfully captures the nearest-neighbor stacking interactions, base-pairing energetics, and environmental bath couplings required to shape a 256-genotype probability distribution. The limitation observed is not biophysical expressivity (the quantum processor’s ability to hold the thermal ensemble), but parameter-space navigation. Just as natural selection struggles to find adaptive peaks on rugged, epistatic fitness landscapes, classical optimizers face rugged parameter landscapes when attempting to reverse-engineer interaction coefficients for multi-nucleotide mutational targets. This demonstrates that variational quantum thermalizers are physically expressive enough to model complex, multi-base mutational hotspots; the frontier lies in refining the search algorithms used to infer those biophysical parameters.

The environmental-coupling sensitivity analysis provides an additional biological interpretation (Fig. 5). Depending on the learned parameter regime, removing the environmental coupling can either increase or decrease the statistical weight of the target sequence. Rather than acting as a simple, unidirectional destabilizer, the environment should be understood as reshaping the effective sequence–energy landscape. This aligns directly with empirical evolutionary genetics: external stressors (such as UV radiation, oxidative conditions, or temperature shifts) do not dictate a single deterministic mutation; instead, they alter the underlying thermodynamic and conformational free energies, stochastically biasing the spectrum of accessible mutational intermediates toward specific hotspots.

From a biological standpoint, our model should be regarded as an effective, coarse-grained statistical description rather than a quantum chemical simulation considering all atoms. Microscopic studies of proton transfer demonstrate that nuclear quantum effects, such as proton tunneling across hydrogen bonds, and environmental bath coupling can modify the reaction barriers and tautomeric stability of individual base-pair configurations [11, 14]. Our framework does not seek to replace detailed quantum chemistry calculations of single base pairs. Instead, it asks whether a coarse-grained Hamiltonian, operating analogously to biophysical nearest-neighbor models in sequence analysis, can aggregate local nucleotide interactions and external stressor couplings into a collective thermal distribution that assigns non-uniform statistical weight to mutation-associated motifs (e.g., mutational hotspots). This builds directly on our previous theoretical framework [15], while the present work contributes direct parameter learning from empirical sequence targets and a matched exact-Gibbs versus VQT benchmark.

This coarse-grained perspective is strongly supported by classical physical biochemistry: electrostatic screening theory predicts that ionic environments systematically alter the free energy of the DNA duplex [25], and single-molecule force spectroscopy confirms sequence-dependent salt corrections to nearest-neighbor base-pair free energies [26]. These established biophysical phenomena validate the core premise of our model: that external abiotic conditions can be effectively captured through coarse-grained energetic coupling terms that reshape the thermodynamic landscape of genotype space.

Mutation bias can strongly orient evolutionary trajectories [2], determining which adaptive paths are traversed by modulating the “arrival of the frequent,” but a static equilibrium target probability is fundamentally distinct from a per-generation mutation rate (*µ*). In vivo, the translation of primary physical changes into heritable substitutions is filtered by active biological machinery: ultraviolet photoproducts are recognized and excised by nucleotide excision repair and transcription-coupled repair pathways [21, 22], while DNA polymerase selectivity, 3^*′*^ → 5^*′*^ exonucleolytic proofreading, and post-replicative mismatch repair introduce additional dynamical rate limits [23, 24]. Direct pedigree sequencing and mutation-accumulation experiments in *Drosophila*, humans, and *Caenorhabditis elegans* yield realized substitution rates at generational timescales [19, 20, 5]. A quantitative mapping connecting the thermodynamic probabilities reported here to those evolutionary rates would therefore require an explicit kinetic framework encompassing environmental exposure duration, lesion repair kinetics, replication fork timing, and ultimately demographic fixation probabilities. We deliberately restrict our scope to thermodynamic fold-enrichment to isolate the raw biophysical inputs before these downstream biological filters act.

The evolutionary interpretation of this work is consequently focused but directly testable. Extensive empirical and theoretical work demonstrates that mutation is not an isotropic, uniform exploration of sequence space [1, 2]; rather, genotype networks are structured by biophysical and biochemical mutational biases, a landscape recently explored using graph-theoretic and quantum-walk formulations [9, 10]. Our results demonstrate that an effective thermal Hamiltonian can provide a first-principles physical mechanism for this structure, generating non-uniform, motif-specific probability distributions over nucleotide strings. The next biological test must be predictive rather than purely reconstructive: interaction parameters should be learned from one empirical set of mutational hotspots (e.g., specific experimental lineages or clinical tumor mutational signatures) and evaluated against held-out flanking sequences, independent mutation-accumulation lines, or orthogonal experimental replicates.

Finally, these findings define a concrete roadmap for expanding the model to longer sequence windows, multi-mode environmental stressors, and cross-validation cohorts. In our benchmarks, the matched comparison with classical theory terminates at *N* = 5 base pairs because exact classical Gibbs calculation requires constructing and exponentiating dense matrices of dimension 2^2*N*+1^ × 2^2*N*+1^, a calculation that exceeded available classical RAM at *N* = 6 (an 8, 192 × 8, 192 dense complex matrix). This steep classical memory wall highlights the practical imperative for a variational quantum formulation, where finite-temperature state preparation and expectation values are evaluated directly via circuit execution and qubit sampling rather than explicit matrix storage. Future implementations can leverage more expressive variational gates, assess finite-measurement sampling noise, incorporate open-system device error mitigation, and execute these simulations on physical quantum processors to scale toward biologically critical sequence motifs (such as promoter elements and transcription factor binding sites) (Fig. 6).

## 5 Conclusions

We have formulated and benchmarked an effective DNA–environment Hamiltonian in which biophysical interaction parameters are learned by maximizing the equilibrium thermodynamic probability of prescribed nucleotide sequences. Exact-Gibbs Bayesian optimization provides the classical quantitative benchmark, enhancing target-sequence probabilities by factors from 9.11 to 159.87 relative to a uniform combinatorial baseline (4^*−N*^ ) across oligomer lengths *N* = 2–5 base pairs.

The Variational Quantum Thermalizer (VQT) addresses this same finite-temperature sequence-bias problem using a parameterized quantum circuit of nine layers. Under the evaluated variational and sampling budgets, VQT successfully preserves target-sequence enrichment above the uniform null expectation across every oligomer length tested. Exact-Gibbs evaluation of the Hamiltonians recovered by the VQT search demonstrates that short sequences accurately reconstruct energy landscapes near the exact classical optimum, whereas larger systems reveal increasing sensitivity to the approximate stochastic nature of nested variational thermal optimization.

Our environmental-coupling sensitivity analysis reveals that interaction with an external stressor selectively reshapes the topology of the sequence-energy landscape, rather than acting as an undifferentiated, global probability amplifier. Taken together, these results establish a direct bridge between a biologically motivated statistical model of DNA–environment thermodynamics and variational quantum algorithms for finite-temperature state preparation.

Beyond establishing algorithmic feasibility, this framework addresses a foundational limitation in evolutionary genetics: the long-standing reliance on isotropic or phenomenological assumptions in sequence-space navigation. For decades, models of genotype networks, adaptive walks, and origin-fixation dynamics have treated mutational transitions as uniform or approximated them via static substitution matrices, largely because computing the true, context-dependent thermodynamic and environmental energetics of DNA across expanding sequence spaces hits an intractable classical memory wall. By demonstrating that a variational quantum circuit can encode and sample environment-coupled thermodynamic biases over discrete nucleotide states, this work provides a first-principles mechanism for how external stressors pre-structure sequence space.

Because a genotype network is an implicitly generated metric graph defined by local physical rules rather than an empirical database, it represents one of the few domains in biology naturally suited to quantum simulation. As quantum hardware scales beyond the reach of classical exact diagonalization (*N* ≥ 6), this approach opens the door to predicting the raw, biophysically accessible mutational spectrum of entire regulatory motifs, promoter architectures, and mutational hotspots under changing climates or chemical exposures. By determining which evolutionary corridors are thermodynamically primed to open—and which remain effectively closed—quantum thermalization offers an analytical foundation for modeling the “arrival of the frequent” from basic physical principles, transforming our ability to anticipate the direction of evolutionary change before natural selection ever intervenes.

## Acknowledgements

HM-D acknowledges the financial support by SECIHTI [CVU 560425]. DS-A acknowledges the financial support of USF via startup funds and the USF Microbiome Research Award 2023. SEVA acknowledges the financial support provided by Tecnologico de Monterrey, Escuela de Ingenieria y Ciencias and SECIHTI-SNII [SNI number 41594]. SEVA acknowledges the unconditional support of his family.

## A Concept translation guide for evolutionary biologists

Here we introduce some of the terms used throughout the manuscript, so that readers unfamiliar with quantum physics can more easily follow the discussion.

### A.1 Statistical-mechanical concepts

### A.2 Variable and operator reference

- *λ*: The parameter vector defining the effective energy model. In the present implementation it contains the coefficients {*h*_*iq*_, *g*_*iq*_, *ϵ*_*iq*_, *ω*} that are optimized to increase the thermal weight of a prescribed target sequence.
- Tr_*b*_(·): Partial trace over the environmental degree of freedom. Since the total system contains both the DNA register and the environment, tracing over the environmental basis produces the reduced DNA state.
- *ρ*_DNA_: The reduced density matrix of the DNA register after the environmental degree of freedom has been traced out. Its diagonal elements define the sequence probabilities used in the analysis.
- ⟨*ψ*_target_|*ρ*_DNA_|*ψ*_target_⟩: Projection onto the prescribed target state. This quantity extracts the target sequence probability *P*_target_ from the full distribution over encoded nucleotide sequences.
- − ln *P*_target_: Negative log-likelihood used as the outer optimization objective. Minimizing this quantity is equivalent to maximizing the thermal probability assigned to the prescribed target sequence.
- ⟨*H*⟩ **and** (Δ*E*)^2^: The mean and variance of the energy in the thermal ensemble. The variance quantifies the magnitude of thermal energy fluctuations of the finite effective model.

### A.3 Variational thermalization concepts

- *F* (*ρ*) = ⟨*H*⟩−*β*^*−*1^*S*(*ρ*): Variational free energy. The VQT minimizes this quantity by balancing low mean energy against the entropy of the trial thermal state.
- *S*(*ρ*) = − Tr(*ρ* log *ρ*): von Neumann entropy. It quantifies how broadly probability weight is distributed over the states of the variational ensemble.
- *p*_*j*_(1) = *σ*(*θ*_*j*_): Bernoulli product distribution used to generate computational-basis input states. Each encoded qubit initially receives an independent probability determined by a logistic function.
- *U*_*ϕ*_|**s**⟩: Parameterized unitary transformation applied to a sampled basis state. The nine-layer circuit rotates and entangles the encoded degrees of freedom, allowing correlations to be represented beyond the initial independent Bernoulli distribution.

#### Nested optimization workflow

The calculation contains two coupled optimization levels. In the *inner loop*, the Hamiltonian is held fixed and the VQT parameters are adjusted to approximate its thermal state by minimizing the variational free energy. In the *outer loop*, the Hamiltonian parameters are varied so as to maximize the probability assigned to the prescribed target sequence.

Accordingly, the inner loop addresses thermal-state preparation, whereas the outer loop addresses Hamiltonian learning.

### A.4 Interpretation of the Hamiltonian parameters

- *ϵ*^⋆^ **versus** *ϵ* = 0: This comparison contrasts the selected environmental coupling with an uncoupled reference. Setting all *ϵ*_*iq*_ = 0 removes the explicit DNA–environment interaction while leaving the remaining Hamiltonian parameters fixed.
- *h*_*iq*_: Local *Z*-field coefficients acting on the two encoded qubit degrees of freedom associated with nucleotide position *i*. They contribute local energetic biases within the effective model.
- *g*_*iq*_: Local transverse *X*-field coefficients acting on the same encoded qubit degrees of freedom. Together with the *Z* terms, they generate noncommuting local dynamics in the effective nucleotide representation.
- *ϵ*_*iq*_: Coupling coefficients between the encoded DNA qubits and the environmental mode. These parameters determine the strength and sign of the explicit environmental contribution to the effective sequence landscape.
- *ω*: Characteristic energy scale of the truncated environmental mode.
- *R*_env_: Environmental sensitivity ratio,

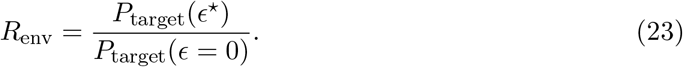

If *R*_env_ = 1, the selected environmental coupling does not change the target probability relative to the uncoupled reference. Values above unity indicate an increase in target statistical weight, whereas values below unity indicate a decrease.

## Notes

### Competing Interest Statement

The authors have declared no competing interest.

